# A high-quality genome of the mangrove *Aegiceras corniculatum* aids investigation of molecular adaptation to intertidal environments

**DOI:** 10.1101/2020.12.28.424522

**Authors:** Xiao Feng, Guohong Li, Shaohua Xu, Weihong Wu, Qipian Chen, Shao Shao, Min Liu, Nan Wang, Cairong Zhong, Ziwen He, Suhua Shi

**Author notes:** These authors contributed equally to this work (Xiao Feng, Guohong Li). Correspondence should be addressed to S.S. or Z.H.

## Abstract

Mangroves have colonized extreme intertidal environments characterized by high salinity, hypoxia, and other abiotic stresses. During millions of years of evolution, mangroves have adapted to these habitats, evolving a series of highly specialized traits. *Aegiceras corniculatum*, a pioneer mangrove species that evolved salt secretion and crypto-vivipary, is an attractive ecological model to investigate molecular mechanisms underlying adaptation to intertidal environments. Here we report a high-quality reference genome of *A. corniculatum* using the PacBio SMRT sequencing technology, comprising 827 Megabases (Mb) and containing 32,092 protein-coding genes. The longest scaffold and N50 for the assembled genome are 13.76 Mb and 3.87 Mb. Comparative and evolutionary analyses revealed that *A. corniculatum* experienced a whole-genome duplication (WGD) event around 35 million years ago after the divergence between *Aegiceras* and *Primula*. We inferred that maintenance of cellular environmental homeostasis is an important adaptive process in *A. corniculatum*. The 14-3-3 protein-coding genes were retained after the recent WGD event, decoding a calcium signal to regulate Na^+^ homeostasis. *A. corniculatum* has more H^+^-ATPase coding genes, essential for the maintenance of low Na^+^ concentration in the cells, than its relatives. Photosynthesis and oxidative-phosphorylation pathways are overrepresented among significantly expanded gene families and might supply the energy needed for salt secretion. Genes involved in natural antioxidant biosynthesis, contributing to scavenging reactive oxygen species against high salinity, have also increased in copy number. We also found that all homologs of *DELAY OF GERMINATION1* (*DOG1*), a pivotal regulator of seed dormancy, lost their heme-binding ability in *A. corniculatum*. This loss may contribute to crypto-vivipary. Our study provides a valuable resource to investigate molecular adaptation to extreme environments in mangroves.

## Introduction

Organismal adaptation is a fundamental topic in evolutionary biology. Intertidal zones are extreme environments that constantly experience drastic changes, with high and fluctuating salinity, hypoxia, and other abiotic stresses (Giri *et al*., 2011). Mangroves have colonized and became well adapted to these habitats, evolving a series of highly specialized traits, including salt tolerance, viviparous embryos, high tannin content, and aerial roots (Ball, 1988a; Liang *et al*., 2008; Parida and Jha, 2010; Tomlinson, 2016). Thus, they are attractive ecological model systems to investigate molecular mechanisms underlying adaptation to intertidal environments.

Vivipary and high salt tolerance are particularly common in mangroves and rare in other woody plants. Salt tolerance is a long-term and dynamic process influenced by multiple genes involving many morphological, physiological, molecular, and cellular processes (Feng *et al*., 2020; Ma *et al*., 2013; Munns and Tester, 2008; Seki *et al*., 2002). Several important signaling pathways have been identified, especially the salt overly sensitive (SOS) signaling pathway, as well as Ca^2+^-dependent and ABA signaling pathways (Ji *et al*., 2013). These systems can maintain and regulate cellular environmental homeostasis through mediating cellular signaling under salt stress. Vivipary in flowering plants is defined as the offspring’s continuous growth when still attached to the mother plant without an apparent dormant period. Embryos of viviparous species, for example, *Rhizophora, Bruguiera, Ceriops*, and *Kandelia* mangroves, first break through the seed coat and then out of the fruit wall. In contrast, crypto-viviparous mangrove species *Aegiceras, Avicennia, Aegialitis*, and *Nypa* only break out of the seed coat but not the fruit wall before dehiscence (Elmqvist and Cox, 1996; Farnsworth, 2000; Tomlinson, 2016; Tomlinson and Cox, 2000). Previous studies have suggested that viviparous propagules are protected from high salinity and other stresses during early development (Liang *et al*., 2008; Zheng *et al*., 1999). These strategies can enhance mangrove reproductive potential in an unstable and extreme environment.

*Aegiceras corniculatum*, one of the most pervasive mangrove species widespread in the Indo West Pacific (IWP) region, is regarded as a pioneer mangrove species for its strong salt tolerance, usually growing along estuary banks. It is a typical outer seaward mangrove (He *et al*., 2019; Tomlinson, 2016). *A. corniculatum* has developed two specialized adaptive traits: salt secretion glands and crypto-vivipary (Ball, 1988b; Deng *et al*., 2009; Ge and Sun, 1999; Parida *et al*., 2004). The lack of omics data for *A. corniculatum* is a major obstacle to understanding the genomics of acquisition and maintenance of these specialized traits and thus this species’ adaptation to intertidal environments. In this study, we report a high-quality reference genome of *A. corniculatum*. Through comparative and evolutionary analyses, we attempt to understand the genomic correlates of phenotypic adaptation to intertidal environments. The genome sequence provided here will accelerate genomic and evolutionary studies of this and other mangrove species.

## Results

### Genome sequencing, assembly, and annotation

We sequenced the genome of *Aegiceras corniculatum*, one of the most pervasive mangrove species widespread in the Indo West Pacific (IWP, Fig. S1), by incorporating PacBio Single-Molecule Real-Time (SMRT) long reads and Illumina short reads. We used a k-mer analysis to estimate genome size at ~ 896 Mb, consistent with the ~841 Mb value obtained by flow cytometry in our previous study (Liu *et al*., 2013; Lyu *et al*., 2018). We generated 139.72 Gb (~169X coverage) of SMRT long reads and 54.71 Gb (~66X coverage) of short reads (Table 1). After preliminary assembly, correction, and polish, the final assembly was 827 Mb, consistent with the estimated genome size. The longest scaffold is 13.76 Mb and N50 is 3.87 Mb (Table1). The GC content of the assembled *A. corniculatum* is 34.37%, similar to closely related species *P. veris* (Nowak *et al*., 2015).

**Table 1.** *A. corniculatum* genome assembly and gene annotation statistics.

| **Genome Features** |  |
| --- | --- |
| **SMRT long reads** | 139.72 Gb |
| **Illumina paired-end reads** | 54.71 Gb |
| **Estimated genome size** | 896 Mb |
| **Assembled genome size** | 826.70 Mb |
| **Scaffold number** | 1494 |
| **N50 length** | 3.87 Mb |
| **N50 count** | 64 |
| **Longest scaffold** | 13.76 Mb |
| **Number of scaffolds ($\geq 1$ Kb)** | 1488 |
| **GC content** | 34.37% |
| **Number of genes** | 32092 |

We examined assembly integrity by aligning CLR subreads to the assembly using minimap2 and Illumina short reads using BWA (Li, 2018; Li and Durbin, 2009). 88.86% of CLR subreads and 97.54 % of Illumina reads were successfully mapped back to our genome. Based on short read mapping, we estimated heterozygosity at ~2.044 sites per kb. We also evaluated the completeness of our genome assembly using BUSCO (Benchmarking Universal Single-Copy Orthologs) v3.1.0 with the eudicotyledons_odb10 lineage dataset (Seppey *et al*., 2019). The BUSCO analysis showed that, at the genomic level, 91.8% of the 2121 expected plant genes were identified as complete, while 6.1% were missing from the assembled *A. corniculatum* genome (Table S1). These results indicate that our genome assembly is of high-quality.

We estimate that 72.10% of the *A. corniculatum* reference genome consists of repetitive sequences. These are predominantly transposable element (TE) families, comprising 560.62 Mb (67.81%) of the genome (Table S2). With repetitive sequences masked, we predicted 32,092 protein-coding genes (Table 1). We were able to functionally annotate 29,207 of them by comparison with public genomic resources from other plants (Table S3). We also predicted 816 tRNAs, 963 rRNAs, 157 snRNAs, and 56 miRNAs (Table S4).

### Phylogenetic position

To determine the position of *A. corniculatum* within Ericales, we reconstructed the phylogeny of 11 angiosperm species with available genome sequences. After searching, aligning, and filtering, we identified 1480 orthologous low-copy genes. Based on these orthologs, we inferred a phylogenetic tree with RAxML-NG using the GTR+GAMMA+I model (Kozlov *et al*., 2019) and *Oryza sativa* (Ouyang *et al*., 2007) as the outgroup (Fig. S2). The phylogenetic relationships among and within the main clades were consistent with previous studies (Chase *et al*., 2016; Yang *et al*., 2020; Zhang *et al*., 2017). We estimate that *Aegiceras* diverged from *Primula* around 42.59 Mya, while Primulaceae diverged from the common ancestor of Ebenaceae, Ericaceae, Actinidiaceae, and Theaceae around 73.81 Mya (Fig. 1).

**Fig. 1.**
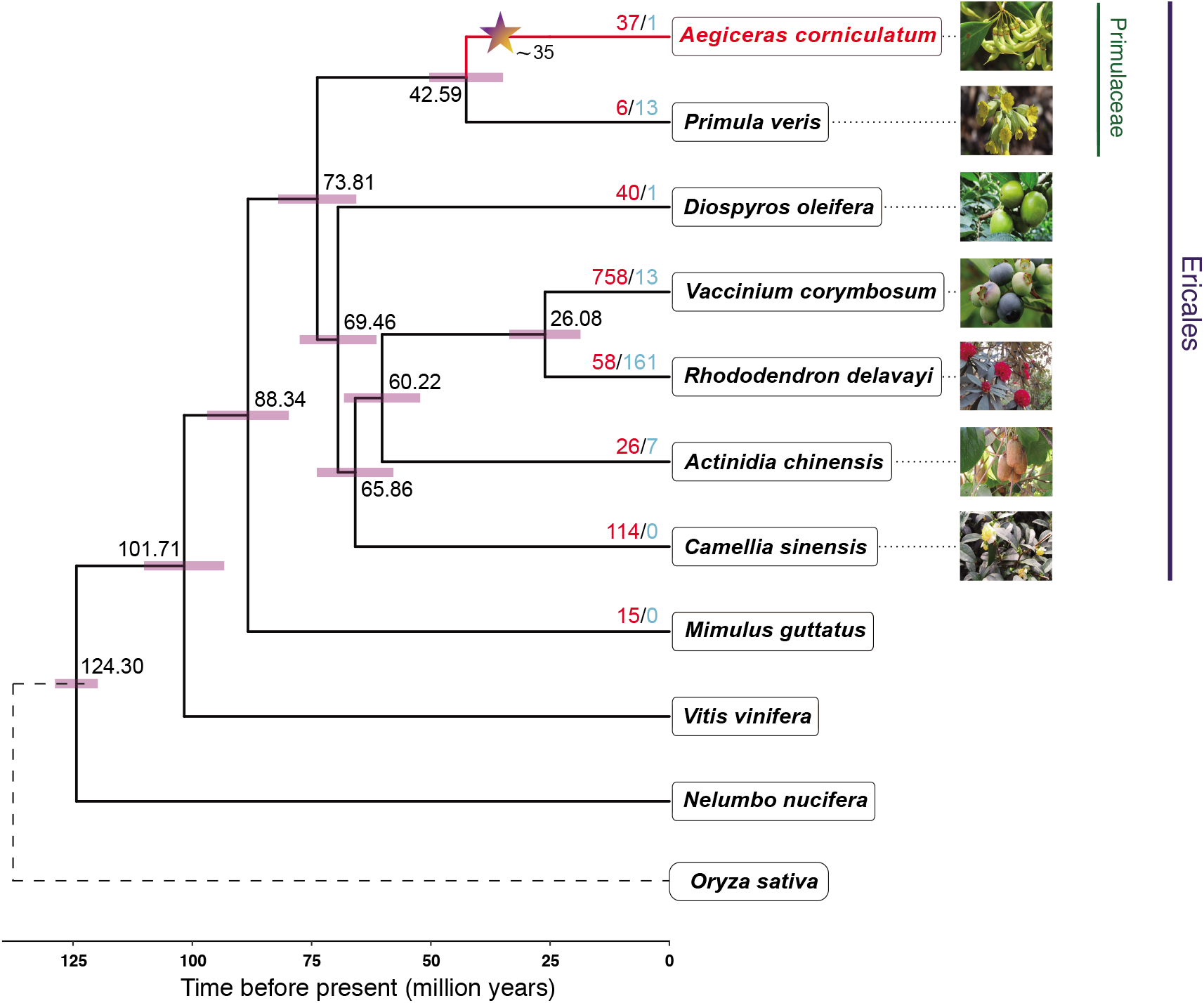
Phylogenetic tree of 11 angiosperm species, including *A. corniculatum* and relatives. Estimated divergence time is shown beside each node. Red bars are 95% confidence intervals. Red and blue numbers indicate significantly expanded and contracted gene families. The dashed line denotes the outgroup *O. sativa*. The star represents the phylogenetic position and estimated date of the *A. corniculatum* lineage-specific WGD event. The credits for morphology pictures are listed in Table S13.

### Gene family analysis

We compared the complexity of gene families between *A. corniculatum* and other species (*P. veris, D. oleifera, V. corymbosum, R. delavayi, A. chinensis, C. sinensis*, and *M. guttatus*) (Colle *et al*., 2019; Hellsten *et al*., 2013; Huang *et al*., 2013; Nowak *et al*., 2015; Suo *et al*., 2020; Xia *et al*., 2019; Zhang *et al*., 2017). We identified 8291 gene families common among these species. Of these, 2663 show signs of expansion in *A. corniculatum*, while another 3096 lost members in this lineage. However, only 37 gene families show statistically significant expansion using CAFE (Mendes *et al*., 2020) in *A. corniculatum* and a single family that contracted since the MRCA of *A. corniculatum* and *P. veris* (Table S5). Gene families encoding proteins involved in ATP-binding cassette transport, oxidative-phosphorylation, and photosynthesis were most obviously expanded (Table S6). The gene families related to natural antioxidant biosynthesis and plant defense also expanded in *A. corniculatum* (Table S7).

### Whole-genome duplication

Whole-genome duplication (WGD) has been widely observed in plants and animals (Clark and Donoghue, 2018; Dehal and Boore, 2005; Van de Peer *et al*., 2017). Polyploid plants may be able to survive and even thrive in extreme environments (He *et al*., 2020; Van De Peer *et al*., 2020; Wu *et al*., 2020). Thus, we wondered whether *Aegiceras* had undergone any recent WGDs. We scanned the genomes of *A. corniculatum* and a closely related species, *P. veris*, using BLASTP and MCScanX. We found that the *A. corniculatum* genome contains 254 syntenic block pairs with at least five shared genes, comprising 6154 (19.18%) protein-coding genes (Fig. S3). This is likely an underestimate of the full extent of genomic duplication, since lost duplicates do not count in this measure. If we include all genes residing within a block delineated by five or more duplicated loci, these blocks cover 18324 (57.10%) genes. In contrast, the *P. veris* genome has only 12 syntenic blocks with 195 (1.07%) genes in them. We also identified 578 such block pairs between the two species. The excess of within-species syntenic block number indicates a possible WGD event within *A. corniculatum* (Fig. 2A). To time this event, we calculated synonymous substitution rates (Ks) between the paralogous genes within the *A. corniculatum* genome. As controls, we looked at Ks distribution between the two species. The mode of the Ks distribution between paralogs within *A. corniculatum* is at a smaller value than for *A. corniculatum vs. P. veris* (Fig. 2B, Fig. S4), reflecting higher similarity and thus indicating that the WGD happened after the divergence between *Aegiceras* and *Primula*. To further estimate the absolute timing of this *A. corniculatum* lineage-specific WGD event, we performed a molecular clock analysis of concatenated gene families (strictly filtered, see the Methods section), calibrated using between-species divergence time. The event was placed at approximately 35 Mya (Fig. 1).

**Fig. 2.**
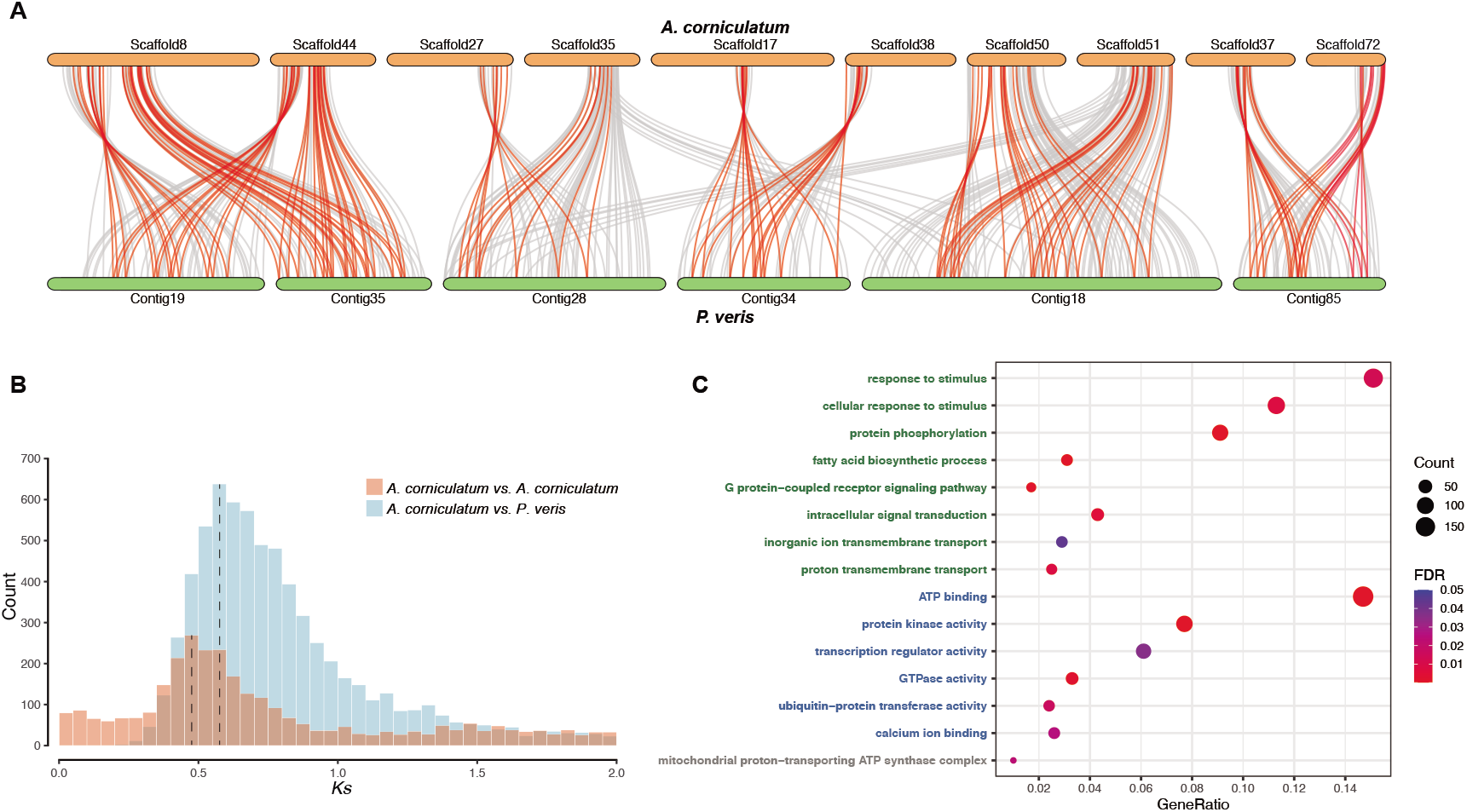
The *A. corniculatum* lineage-specific whole-genome duplication (WGD) event. (**A**) Diagram of syntenic blocks between genomic regions from *A. corniculatum* and the closely related species *P. veris*. Red links represent collinear gene pairs with a 1:2 ratio in the two species. (**B**) Ks distribution between paralogous genes within the same species and orthologous genes between the two species. Dashed lines denote the WGD event (left) and speciation event (right). (**C**) Gene Ontology enrichment among retained gene duplicates compared with single genes. The size and color of the bubbles represent gene number and FDR value calibrated using the Benjamini-Hochberg method. The green, blue, and grey categories stand for the Biological Process, Molecular Function, and Cellular Component, respectively.

Genes acquired during a large-scale duplication event are typically rapidly lost. Retained duplicates are potentially enriched for genes whose increased expression or acquired functions aid survival in the newly acquired habitat (Freeling, 2009; Van de Peer *et al*., 2017). With single-copy genes as a control, we performed GO enrichment and pathway enrichment analyses on the genes retained after the *Aegiceras*-specific WGD. We found that most of the genes were involved in response to stimulus, protein phosphorylation, inorganic ion transmembrane transport, ATP binding, protein kinase activity, transcription regulator activity, calcium ion binding, and mitochondrial proton-transporting ATP synthase complex GO categories. The NOD-like-receptor signaling, MAPK signaling, calcium signaling, and plant hormone signal transduction pathways were also overrepresented (Fig. 2C, Table S8). The preferential retentions of transcription regulation, signal transduction, and energy metabolism pertain to the adaptation to intertidal zones.

### High-salt adaptation-supportive mechanisms

The most extreme challenge for species living in intertidal zones is the unstable and high salinity due to constant tidal fluctuations. *A. corniculatum* has evolved typical salt glands on leaves to enhance its salt tolerance. Among the genes retained after the recent WGD, the calcium-activated 14-3-3 protein-coding gene is particularly interesting as it is a molecular switch in salt stress tolerance, decoding a calcium signal to enhance plant salt tolerance (Yang *et al*., 2019) (Fig. 3A). After a strict alignment with the Pfam and NR databases, and the *A. thaliana* annotation, we identified fourteen 14-3-3 protein-coding genes in *A. corniculatum*, more than the nine in *P. veris* (Fig. 3B, Table S9). We also identified 19 H^+^-ATPase coding genes in *A. corniculatum*, more than in its relatives (Fig. 3C, Table S10). After the recent WGD event, twelve duplicates of 14-3-3 protein-coding genes and four duplicates of H^+^-ATPase coding genes were retained. We note that many retentions were annotated with functions involving calcium signal-activated salt overly sensitive (SOS) pathway, in particular *J3* and calcineurin related genes. This result confirms the GO enrichment and pathway enrichment analyses and shows preferential recent WGD retentions of signal transduction and energy metabolism during salt adaptation.

**Fig. 3.**
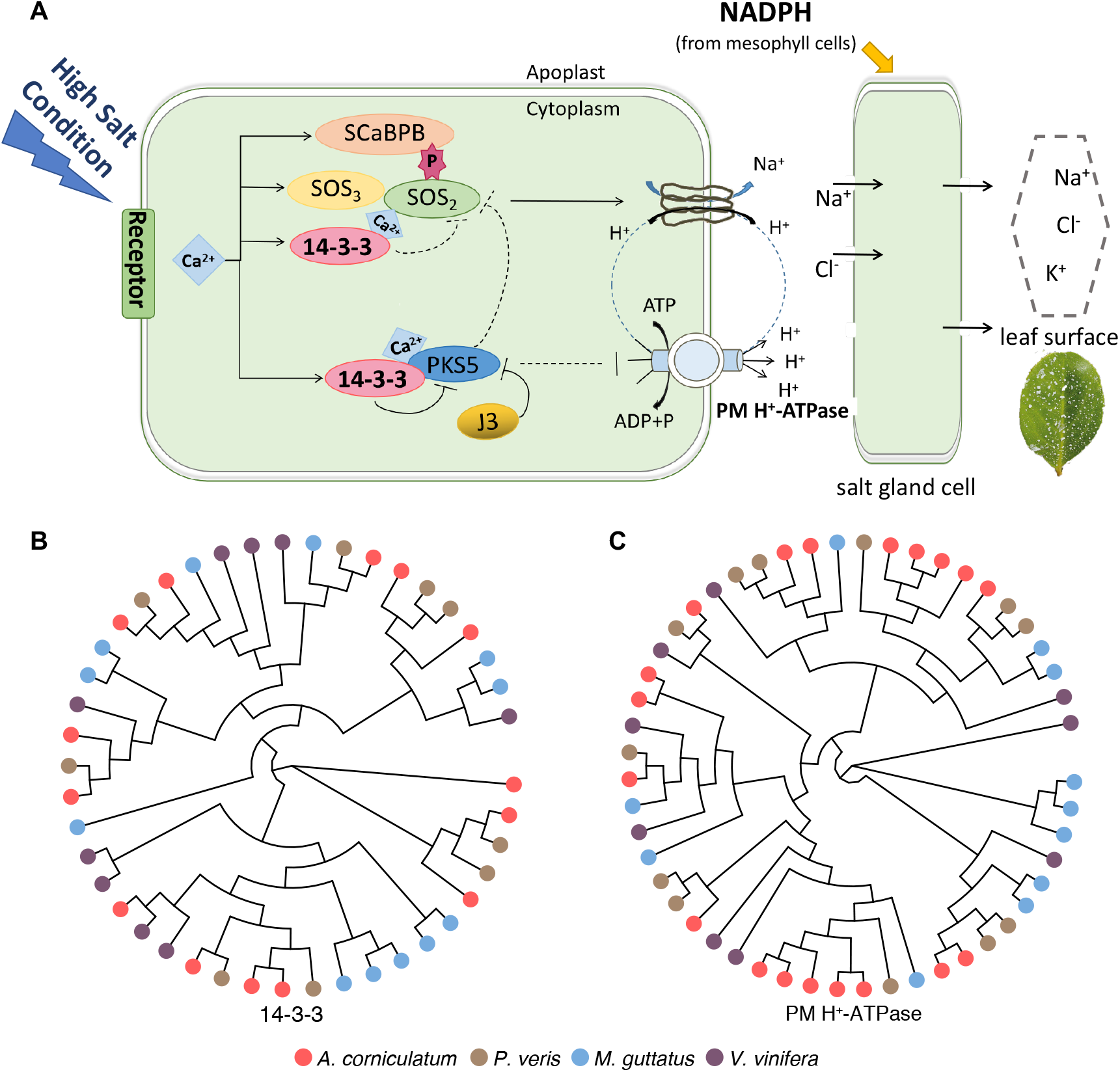
The possible pathway of Na^+^ transport and salt secretion in *A. corniculatum*. **(A)** The 14-3-3 protein-coding gene, decoding a calcium signal to enhance plant salt tolerance, can activate the SOS pathway under high salinity conditions. Na^+^ is transported into the salt gland and then secreted onto the leaf surface. Modified from Yuan *et al*., 2016 and Yang *et al*., 2019. Unrooted phylogenetic tree of 14-3-3 **(B)** and PM H+-ATPase **(C)** protein-coding genes in *A. corniculatum* and relatives.

### Identification of a pivotal gene loss accounting for crypto-vivipary

In order to explore genetic mechanisms underlying the emergence of crypto-vivipary in *A. corniculatum*, we focused on homologs of *DELAY OF GERMINATION1* (*DOG1*), a pivotal regulator controlling seed dormancy (Nishimura *et al*., 2018; Nonogaki, 2019). We searched for putative homologs of *AtDOG1* in each Ericales genome and selected the best hits. Despite the broad sequence diversity of *DOG1* (Table S12), we discovered a conspicuous amino acid divergence that appears exclusively in one *A. corniculatum* homolog Aco04515 within a relatively conserved region at position 633 of the multi-sequence alignment. All other genes have a histidine residue at this position while it is replaced by glutamine in Aco04515 of *A. corniculatum* (Fig. S6A). This residue, His^245^ of AtDOG1, is critical heme-binding site formation. Together with His^249^, His^245^ serves as an axial ligand for the heme (Fig. 4A). This has been demonstrated in a transgenic experiment that heme-binding via His^245^ and His^249^ (Nishimura *et al*., 2018).

**Fig. 4.**
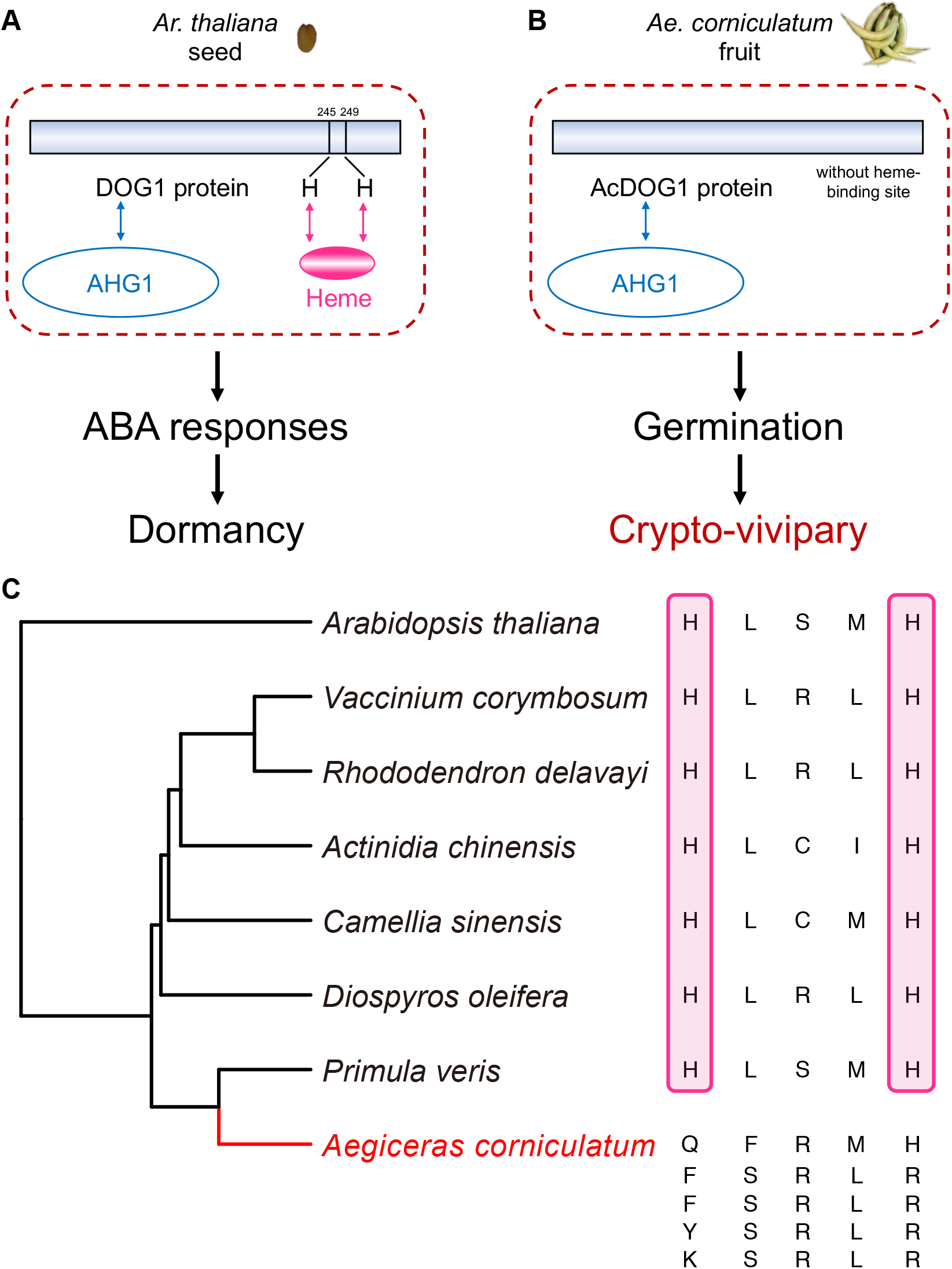
All DOG1 homologs lost the ability to bind heme in *A. corniculatum* leading to functional deficiency. (**A**) Mechanism of seed dormancy induced by AtDOG1 with the heme-binding site. Modified from Nishimura *et al*., 2018 and Nonogaki, 2019. (**B**) Loss of the heme-binding site in the homologous proteins prompts seed germination, probably accounting for the specialized crypto-vivipary trait. (**C**) Amino acid alignment of the heme-binding site among AtDOG1 proteins, putative homologs in each Ericales genome. The red frame highlights the two critical histidine residues of the heme-binding site.

We inferred that the incompleteness of the heme-binding site caused by missing one of the two histidine residues might result in a functional deficiency of *DOG1* homologs in *A. corniculatum* (Fig. 4B). Considering the genetic divergence of *DOG1* among Asterids plant species (Table S11, Table S12, Fig. S5) and the abundance of *DOG1-like* homologs in *A. corniculatum*, we wondered whether there is any potential compensation achieved by other *DOG1* or *DOG1-like* homologs with a complete heme-binding site. Thus, we searched for all potential candidate *DOG1* genes (see Methods), but found that all candidates appeared to have lost critical histidine residues (Fig. 4C). For each candidate, we further searched for the best hit in *A. thaliana* as well as other Ericales genomes (Supplementary Fig. S6B-F). To exclude the possibility of assembly failure, we mapped more accurate Illumina short reads we generated for *A. corniculatum* to the *DOG1* region of two closely related species, *P. veris* and *C. sinensis*. These alignments show the same loss of critical His amino acids. These results indicate no functional heme-binding *DOG1* homolog exists in *A. corniculatum*.

## Discussion

As a pioneer mangrove species in the Primulaceae family, *A. corniculatum* has a series of specialized adaptive traits, including salt secretion and crypto-vivipary. A combination of Ks-based, synteny, and molecular clock analyses shows that *A. corniculatum* experienced a recent round of WGD since the well-known paleo-hexaploidization (γ WGD) event shared by core eudicots (Jiao *et al*., 2012). Genes preferentially retained after this duplication are disproportionately involved in stimulus response, ion transport, signal transduction, and energy metabolism. The evidence presented here suggests that the lineage-specific WGD event might have contributed to the adaptation of *A. corniculatum* to the fluctuating high salinity of tropical intertidal environments.

In general, salt adaptation is both a long-term and dynamic process and involves many morphological, physiological, cellular, and molecular processes (Feng *et al*., 2020; Ma *et al*., 2013; Munns and Tester, 2008; Seki *et al*., 2002). Mangrove trees can induce specific salt-responsive genes and transcription factors to facilitate adaptation to intertidal environments (Feng *et al*., 2020). The salt overly sensitive (SOS) pathway is a well-defined signaling pathway for controlling ion homeostasis at the cellular and tissue level (Ji *et al*., 2013). The 14-3-3 protein decoding salt-induced calcium signals can activate SOS1 (plasma membrane Na^+^/H^+^ antiporter) and PM H^+^-ATPase to adapt to salt stress (Yang *et al*., 2019). Our comparative genomic analysis reveals that 14-3-3 and H^+^-ATPase protein-coding genes have more copies in *A. corniculatum* than in its relatives (*P. veris, M. guttatus*, and *V. vinifera*).

Salt secretion via glands is a highly specialized trait in mangroves to enhance their salt tolerance (Shi *et al*., 2005). This mechanism involves an increase of H^+^-ATPase expression as salt secretion accelerates in high-salt environments (Chen *et al*., 2010). Salt secretion places high energy demands on the plant. Mesophyll cells can provide photoassimilates and NADPH (the sources of energy for salt excretion) to the salt gland cells via plasmodesmata (Yuan *et al*., 2016). In this context, it is noteworthy that photosynthesis and oxidative-phosphorylation pathway are expanded in the *A. corniculatum* genome.

The biggest challenge for mangrove trees surviving in the extreme and harsh intertidal environment is the unstable and strong ecological pressure. Our findings suggest that the maintenance of cellular environmental homeostasis is an important adaptive process in *A. corniculatum*. Salt ions are transported from mesophyll cells into the salt gland and then secreted (Fig. 3A). Expansion of antioxidant biosynthesis related genes can contribute to scavenging reactive oxygen species (ROS) against hypersaline stress. The functional enrichment of genes retained after WGD also supports homeostasis maintenance as an important adaptation driver.

Unlike regular seed development, vivipary or crypto-vivipary in mangrove seeds can decrease dormancy and promote germination when still attached to the mother plant. It is now well established that *DOG1* is a key regulator controlling seed maturation, dormancy, and germination in plants (Dekkers *et al*., 2016; Nakabayashi *et al*., 2012; Nonogaki, 2019). *DOG1* belongs to a small gene family consisting of *DOG1* and four additional *DOG1-like* genes. They are found in many species, such as rice, wheat, barley, sorghum, and Brassicaceae including *A. thaliana* (Ashikawa *et al*., 2010, 2013; Bentsink *et al*., 2006; Carrillo-Barral *et al*., 2015; Graeber *et al*., 2010; Sugimoto *et al*., 2010). Since *DOG1* was first identified and cloned in *A. thaliana* as a major quantitative trait locus for seed dormancy (Bentsink *et al*., 2006), most mechanisms of seed dormancy have been elucidated in this species (Nonogaki, 2019). Recent genomic analyses in mangrove trees suggest that amino acid substitution and positive selection of essential seed development genes, expression changes of key plant hormone metabolism and embryonic genes, reduction of proanthocyanidin and storage protein production, and *DOG1* loss together contribute to vivipary in the Rhizophoraceae family (Qiao *et al*., 2020; Xu *et al*., 2017). In the crypto-viviparous mangrove *A. corniculatum* embryos only break out of the seed coat before dehiscence. Considering the genetic divergence of *DOG1* among Asterids plant species (Table S11, Table S12) and the wide distribution of functional *DOG1-like* genes in cereals (Ashikawa *et al*., 2013; Graeber *et al*., 2014), we searched all potential *DOG1* or *DOG1-like* homologs in *A. corniculatum* and its relatives. The complete heme-binding site found in other species appears to have been lost in *A. corniculatum* (Fig. 4C). Previous studies revealed that heme binding at His^245^ and His^249^ of DOG1 is critical to its seed dormancy function (Nishimura *et al*., 2018). Thus, losing the complete heme-binding site promotes seed germination, probably enabling crypto-vivipary (Fig. 4B), protecting propagules from stresses during early development. This strategy can enhance *A. corniculatum*’s fertility in an unstable and extreme environment.

In summary, we constructed a high-quality genome assembly for *Aegiceras corniculatum* by combining SMRT long reads with highly accurate short reads. It provides new insights into the evolution of two specialized adaptive traits (salt secretion and crypto-vivipary) in mangroves. We deduce that the maintenance of cellular environmental homeostasis is an important adaptive process. Our study also will be helpful for genetic, genomic, and evolutionary studies in both *Aegiceras* and other plants inhabiting extreme environments.

## Methods

### Plant material

We sampled one mature *Aegiceras corniculatum* plant from the nursery of Dongzhai Harbor National Nature Reserve in Hainan with permission. Fresh and healthy leaves were harvested and immediately frozen in liquid nitrogen, followed by preservation at −80 °C in the laboratory before DNA extraction. High-quality genomic DNA was extracted from leaves using the modified CTAB method. RNase A was used to remove RNA contaminants. The quality of the extracted DNA was examined using a NanoDrop 2000 spectrophotometer (NanoDrop Technologies, Wilmington, DE, USA). The quantity was estimated using electrophoresis on a 0.8% agarose gel. Total RNA from leaves was extracted using the TRIzol reagent (Invitrogen) according to the manufacturer’s protocol.

### PacBio long-read library preparation and sequencing

Single-Molecule Real-Time (SMRT) long read sequencing was performed on a PacBio sequel II platform (Pacific Biosciences, Menlo Park, CA, USA). A single SMRT-bell library with 40 kb long inserts was constructed from sheared genomic DNA using a template library preparation workflow. The library was sequenced on PacBio SMRT cells 8M (acquiring one movie of 15 hours per SMRT cell) using a PacBio Sequel II instrument. After data filtering and preprocessing, 10.24 million long reads were generated, yielding ~139.72 Gb (169X coverage) with an average read length of 13,647 bp.

### Short-read sequencing

For DNA short-read sequencing, 150 bp paired-end libraries were prepared for sequencing on Illumina NovaSeq 6000 platform and yielded ~54.71 Gb of bases. The RNA-seq library was sequenced on BGI-seq 500 platform for gene prediction and yielded ~8.06 Gb of bases.

### *De novo* genome assembly

We assembled the *A. corniculatum de novo* genome based on the PacBio long reads using three assemblers: Canu (Koren *et al*., 2017), Falcon (Chin *et al*., 2016), and Mecat2 (Xiao *et al*., 2017) with optimized parameters. The optimal assembly (using Mecat2) was further polished with racon (v.1.3.1) using long reads (Vaser *et al*., 2017). To improve primary assembly accuracy, we corrected the remaining errors using pilon (v1.22) based on Illumina short reads (Walker *et al*., 2014). Purge Haplotigs was used to filter redundant sequences due to heterozygosity (Roach *et al*., 2018).

### Genome annotations

We identified repetitive sequences in the *A. corniculatum* genome by integrating homology-based and *de novo* approaches. For homology-based prediction, we identified the known TEs within the *A. corniculatum* genome using RepeatMasker (open-4.0.9) with the Repbase TE library (Bao et al., 2015; Tarailo-Graovac and Chen, 2009). RepeatProteinMask searches were also conducted using the TE protein database as a query library. For *de novo* prediction, we constructed a *de novo* repeat library of the *A. corniculatum* genome using RepeatModeler (Flynn *et al*., 2020) to comprehensively conduct, refine, and classify consensus models of putative interspersed repeats for the *A. corniculatum* genome. Furthermore, we performed a *de novo* search for long terminal repeat (LTR) retrotransposons against the *A. corniculatum* genome sequences using LTR_FINDER (v1.0.7) (Xu and Wang, 2007). We also identified tandem repeats using the Tandem Repeat Finder (TRF) package (Benson, 1999) and the non-interspersed repeat sequences, including low-complexity repeats, satellites, and simple repeats, using RepeatMasker. Finally, we merged these library files of the two methods and used RepeatMasker to identify the repeat contents.

We conducted annotation of protein-coding genes in the *A. corniculatum* genome using a combination of *de novo*, homology-based, and RNA-seq-based prediction. We used Augustus (v3.3.1) (Stanke *et al*., 2006) and GlimmerHMM (Majoros *et al*., 2004) to perform *de novo* gene prediction based on the masked genome. Protein sets were collected and chosen as homology-based evidence from sequenced plants Ericales (*Actinidia chinensis* (Huang *et al*., 2013), *Camellia sinensis* (Xia *et al*., 2019), *Diospyros oleifera* (Suo *et al*., 2020), *Primula veris* (Nowak *et al*., 2015), *Rhododendron delavayi* (Zhang *et al*., 2017), *Vaccinium corymbosum* (Colle *et al*., 2019)), and model plants (*Arabidopsis thaliana* (Lamesch *et al*., 2012) and *Oryza sativa* (Ouyang *et al*., 2007)). Clean RNA-seq reads were used as RNA-seq-based evidence; gene structure was estimated using Tophat2 (v2.1.1) and Cufflinks (v2.2.1) (Kim *et al*., 2013; Langmead and Salzberg, 2012; Trapnell *et al*., 2012). Finally, Maker (v3.00) was used to integrate all evidence to generate non-redundant gene models (Cantarel *et al*., 2007). Functional annotations were assigned according to the best match of the alignments to the NCBI (NR), Swissprot, TrEMBL, InterPro, the Kyoto Encyclopedia of Genes and Genomes (KEGG), Gene Ontology (GO), and Pfam non-redundant protein databases.

### Genome quality assessment

The quality-filtered reads were used for genome size estimation. First, we generated a 17-mer occurrence distribution of sequencing reads from short libraries using the k-mer method. Then, we estimated the genome size using GCE (v.1.0.2) (Liu *et al*., 2013). Flow cytometry analysis for the measurement of nuclear DNA content was performed in our previous study (Lyu *et al*., 2018). The genome size of *A. corniculatum* is approximately 841 Mb (flow cytometry) or 896 Mb (k-mer analysis). In order to examine the assembly integrity, the Continuous Long Reads (CLR) subreads were aligned to the assembly using minimap2 (v2.5) (Li, 2018), and the Illumina short reads were also aligned to the assembly using BWA (Li and Durbin, 2009). We evaluated the completeness of the assembly and gene prediction using BUSCO v3.1.0 based on the eudicotyledons_odb10 database (2121 single-copy genes) (Seppey *et al*., 2019).

### Phylogenetic analyses

We used OrthoFinder (Emms and Kelly, 2019) to identify orthologous genes from *A. corniculatum* and ten other angiosperms species: *Primula veris* (Nowak *et al*., 2015), *Diospyros oleifera* (Suo *et al*., 2020), *Vaccinium corymbosum* (Colle *et al*., 2019), *Rhododendron delavayi* (Zhang *et al*., 2017), *Actinidia chinensis* (Huang *et al*., 2013), *Camellia sinensis* (Xia *et al*., 2019), *Mimulus guttatus* (Hellsten *et al*., 2013), *Vitis vinifera* (Jaillon *et al*., 2007), *Nelumbo nucifera* (Shi *et al*., 2020), *Oryza sativa* (Ouyang *et al*., 2007). We also used the reciprocal BLASTP best-hit method for the 11 angiosperm proteins and single-copy genes from the BUSCO dataset. We then merged these two results to identify low-copy genes. We individually aligned low-copy orthologous proteins using MAFFT (Katoh, 2002; Katoh and Standley, 2013) and used the aligned protein sequences to generate codon alignments using PAL2NAL (Suyama *et al*., 2006). We further trimmed alignments using Gblocks 0.91b (Castresana, 2000) and discarded alignments shorter than 150 bp. Based on these alignments, we inferred a phylogenetic tree using RAxML-NG with the GTR+GAMMA+I model, 1000 bootstrap replicates (Kozlov *et al*., 2019), and *Oryza sativa* as an outgroup (Ouyang *et al*., 2007). Following its reconstruction, we further dated the tree using MCMCTREE from the PAML package with approximate likelihood calculation (Yang, 2007). This method provides a fast and efficient way of analyzing large datasets using complex models (Reis and Yang, 2011). Two reliable fossil calibrations were incorporated (Morris *et al*., 2018). First, the root node of eudicots and monocots was constrained between 125-247 Myr before present. Second, the common ancestor of eudicots was placed at 119.6-128.63 Myr before present. The MCMC analyses were run for 10 million generations and sampled every 500 generations after a burn-in of 1000000 iterations. The MCMC analyses were run twice independently to ensure convergence. We used the R package ggtree to visualize the phylogenetic tree (Yu *et al*., 2017). The origin and credit for the plant pictures are listed in Table S13.

Gene family evolution in Ericales was examined using CAFE (Mendes *et al*., 2020). We obtained counts of gene families and genes in seven species of Ericales and *M. guttatus* from OrthoFinder (Emms and Kelly, 2019) and removed the large gene families with more than 100 gene copies in one or more species. We identified gene family expansions or contractions only when the gene count change was significant with a *P*-value < 0.01. According to *A. corniculatum* genome annotations, we assigned these gene families to KEGG pathways and compared these pathways’ gene numbers in the expanded families and the whole genome.

### Whole-genome duplication analyses

To detect the degree of collinearity, we aligned protein sequences between *A. corniculatum* and the more closely related species *P. veris* and within each species using BLASTP (with identity ≥ 30%, e-value < 1E-10, alignment length ≥ 30% of both query and reference sequences). We then identified syntenic blocks containing a minimum of five shared genes using MCScanX (Wang *et al*., 2012). Syntenic blocks were visualized by TBtools (Chen *et al*., 2020) and Circos (Krzywinski *et al*., 2009). For analyses of the WGD events, we first obtained alignments of all gene pairs as described above. Then we used KaKs_Calculator to calculate synonymous substitution rates (Ks) with the YN substitution model and visualize the Ks distributions for paralogous genes within *A. corniculatum* and orthologous genes between *A. corniculatum* and *P. veris* (Wang *et al*., 2010). To identify paralogous genes generated in the lineage-specific WGD on the *A. corniculatum* branch, we filtered all gene pairs except those with Ks in the 0.2~1.1 range for further analysis. We also identified single genes using the duplicate_gene_classifier module from MCScanX (Wang *et al*., 2012). With single genes as a control, we performed GO and KEGG enrichment analyses of all filtered paired genes belonging to collinear blocks using BiNGO in Cytoscape_v3.7.2 (Shannon *et al*., 2003).

To gauge the date of the *A. corniculatum* lineage-specific WGD event, we estimated the WGD event’s absolute timing through molecular clock analysis of concatenated gene families, calibrated using species divergence times, similarly to the approach adopted by Clark and Donoghue (Clark and Donoghue, 2017). Based on gene duplicates generated by the WGD, we first identified *A. corniculatum vs. P. veris* orthogroups that were at exactly 2:1 copy number ratio according to collinear relationships identified above. The corresponding orthologs from *R. delavayi* and *V. vinifera* in each orthogroup were identified by searching for the *P. veris* gene’s best hit. To further verify a clear signal of the recent WGD event in the selected gene families, we built individual gene trees based on multiple sequence alignments of these four plant species as described above and performed gene tree reconstructions using RAxML-NG (Kozlov *et al*., 2019). We discarded orthogroups whose topology did not clearly reflect the WGD event signal or was incongruent with the species tree. Of the 552 retained orthogroups, 136 gene families had a clear signal of the lineage-specific WGD event on the *A. corniculatum* branch. To improve the robustness and precision of estimation, we concatenated 136 multiple sequence alignments with a random combination of each two gene copies in *A. corniculatum*. The nodes were constrained using species divergence times from the phylogenetic tree described above. Molecular clock analysis was conducted on concatenated alignments using the approximate likelihood calculation method in MCMCTREE under the appropriate model (Reis and Yang, 2011; Yang, 2007). We also reconstructed the topology based on our concatenated alignments using RAxML-NG and found it to coincide exactly with the constrained tree. Each analysis was run twice independently to ensure convergence.

### Identification of the key crypto-vivipary gene loss

We searched for candidate homologs of *DELAY OF GERMINATION1* (*DOG1*) in all seven Ericales genomes against *A. thaliana* genome data (TAIR10, www.arabidopsis.org) using BLASTP. We searched for *A. corniculatum* candidate homologs of both *DOG1* and *DOG1-like* genes and retained all hits with an e-value cutoff of 1E-10. Only the best hits were retained for further analysis from the other six species. We aligned all the homologs together to verify their heme-binding potential. To identify the exclusive ortholog of each *A. corniculatum* candidate in *A. thaliana* as well as other Ericales species, we performed BLASTP searches using *A. corniculatum* as a database and selected the best hit. All alignments mentioned above were implemented using the accurate option (L-INS-i) of MAFFT (Katoh, 2002; Katoh and Standley, 2013). Alignment results were displayed using AliView (Larsson, 2014). Considering the heme-binding site is essential for the *DOG1* function, we mapped Illumina short-reads of *A. corniculatum* to the potential *DOG1* region of two relatives, *P. veris* and *C. sinensis*, and manually checked the alignments. To estimate *DOG1* genetic divergence, we employed the Kimura two-parameter method (Kimura, 1980) to calculate the genetic divergence among 21 Asterids and three outgroups (Table S11, Fig. S5).

## References

Ashikawa, I., Abe, F., and Nakamura, S. (2013) DOG1-like genes in cereals: Investigation of their function by means of ectopic expression in Arabidopsis. Plant Sci., 208, 1–9.

Ashikawa, I., Abe, F., and Nakamura, S. (2010) Ectopic expression of wheat and barley DOG1-like genes promotes seed dormancy in Arabidopsis. Plant Sci., 179, 536–542.

Ball, M.C. (1988a) Ecophysiology of mangroves. Trees, 2, 129–142.

Ball, M.C. (1988b) Salinity Tolerance in the Mangroves Aegiceras corniculatum and Avicennia marina. I. Water Use in Relation to Growth, Carbon Partitioning, and Salt Balance. Funct. Plant Biol., 15, 447–464.

Bao, W., Kojima, K.K., and Kohany, O. (2015) Repbase Update, a database of repetitive elements in eukaryotic genomes. Mob. DNA, 6, 11.

Benson, G. (1999) Tandem repeats finder: a program to analyze DNA sequences. Nucleic Acids Res., 27, 573–580.

Bentsink, L., Jowett, J., Hanhart, C.J., and Koornneef, M. (2006) Cloning of DOG1, a quantitative trait locus controlling seed dormancy in Arabidopsis. Proc. Natl. Acad. Sci., 103, 17042–17047.

Cantarel, B.L., Korf, I., Robb, S.M.C., Parra, G., Ross, E., Moore, B., et al. (2007) MAKER: An easy-to-use annotation pipeline designed for emerging model organism genomes. Genome Res., 18, 188–196.

Carrillo-Barral, N., Matilla, A.J., García-Ramas, C., and Rodríguez-Gacio, M. del C. (2015) ABA-stimulated SoDOG1 expression is after-ripening inhibited during early imbibition of germinating Sisymbrium officinale seeds. Physiol. Plant., 155, 457–471.

Castresana, J. (2000) Selection of Conserved Blocks from Multiple Alignments for Their Use in Phylogenetic Analysis. Mol. Biol. Evol., 17, 540–552.

Chase, M.W., Christenhusz, M.J.M., Fay, M.F., Byng, J.W., Judd, W.S., Soltis, D.E., et al. (2016) An update of the Angiosperm Phylogeny Group classification for the orders and families of flowering plants: APG IV. Bot. J. Linn. Soc., 181, 1–20.

Chen, C., Chen, H., Zhang, Y., Thomas, H.R., Frank, M.H., He, Y., and Xia, R. (2020) TBtools: An Integrative Toolkit Developed for Interactive Analyses of Big Biological Data. Mol. Plant, 13, 1194–1202.

Chen, J., Xiao, Q., Wu, F., Dong, X., He, J., Pei, Z., and Zheng, H. (2010) Nitric oxide enhances salt secretion and Na+ sequestration in a mangrove plant, Avicennia marina, through increasing the expression of H+-ATPase and Na+/H+ antiporter under high salinity. Tree Physiol., 30, 1570–1585.

Chin, C.-S., Peluso, P., Sedlazeck, F.J., Nattestad, M., Concepcion, G.T., Clum, A., et al. (2016) Phased diploid genome assembly with single-molecule real-time sequencing. Nat. Methods, 13, 1050–1054.

Clark, J.W. and Donoghue, P.C.J. (2017) Constraining the timing of whole genome duplication in plant evolutionary history. Proc. R. Soc. B Biol. Sci., 284, 20170912.

Clark, J.W. and Donoghue, P.C.J. (2018) Whole-Genome Duplication and Plant Macroevolution. Trends Plant Sci., 23, 933–945.

Colle, M., Leisner, C.P., Wai, C.M., Ou, S., Bird, K.A., Wang, J., et al. (2019) Haplotype-phased genome and evolution of phytonutrient pathways of tetraploid blueberry. Gigascience, 8, 1–15.

Dehal, P. and Boore, J.L. (2005) Two Rounds of Whole Genome Duplication in the Ancestral Vertebrate. PLoS Biol., 3, e314.

Dekkers, B.J.W., He, H., Hanson, J., Willems, L.A.J., Jamar, D.C.L., Cueff, G., et al. (2016) The Arabidopsis DELAY OF GERMINATION 1 gene affects ABSCISIC ACID INSENSITIVE 5 (ABI5) expression and genetically interacts with ABI3 during Arabidopsis seed development. Plant J., 85, 451–465.

Deng, S., Huang, Y., He, H., Tan, F., Ni, X., Jayatissa, L.P., et al. (2009) Genetic diversity of Aegiceras corniculatum (Myrsinaceae) revealed by amplified fragment length polymorphism (AFLP). Aquat. Bot., 90, 275–281.

Elmqvist, T. and Cox, P.A. (1996) The Evolution of Vivipary in Flowering Plants. Oikos, 77, 3–9.

Emms, D.M. and Kelly, S. (2019) OrthoFinder: phylogenetic orthology inference for comparative genomics. Genome Biol., 20, 238.

Farnsworth, E. (2000) The Ecology and Physiology of Viviparous and Recalcitrant Seeds. Annu. Rev. Ecol. Syst., 31, 107–138.

Feng, X., Xu, S., Li, J., Yang, Y., Chen, Q., Lyu, H., et al. (2020) Molecular adaptation to salinity fluctuation in tropical intertidal environments of a mangrove tree Sonneratia alba. BMC Plant Biol., 20, 178.

Flynn, J.M., Hubley, R., Goubert, C., Rosen, J., Clark, A.G., Feschotte, C., and Smit, A.F. (2020) RepeatModeler2 for automated genomic discovery of transposable element families. Proc. Natl. Acad. Sci., 117, 9451–9457.

Freeling, M. (2009) Bias in Plant Gene Content Following Different Sorts of Duplication: Tandem, Whole-Genome, Segmental, or by Transposition. Annu. Rev. Plant Biol., 60, 433–453.

Ge, X.J. and Sun, M. (1999) Reproductive biology and genetic diversity of a cryptoviviparous mangrove Aegiceras corniculatum (Myrsinaceae) using allozyme and intersimple sequence repeat (ISSR) analysis. Mol. Ecol., 8, 2061–2069.

Giri, C., Ochieng, E., Tieszen, L.L., Zhu, Z., Singh, A., Loveland, T., et al. (2011) Status and distribution of mangrove forests of the world using earth observation satellite data. Glob. Ecol. Biogeogr., 20, 154–159.

Graeber, K., Linkies, A., Müller, K., Wunchova, A., Rott, A., and Leubner-Metzger, G. (2010) Cross-species approaches to seed dormancy and germination: conservation and biodiversity of ABA-regulated mechanisms and the Brassicaceae DOG1 genes. Plant Mol. Biol., 73, 67–87.

Graeber, K., Linkies, A., Steinbrecher, T., Mummenhoff, K., Tarkowska, D., Ture kova, V., et al. (2014) DELAY OF GERMINATION 1 mediates a conserved coat-dormancy mechanism for the temperature- and gibberellin-dependent control of seed germination. Proc. Natl. Acad. Sci., 111, E3571–E3580.

He, Z., Li, X., Yang, M., Wang, X., Zhong, C., Duke, N.C., et al. (2019) Speciation with gene flow via cycles of isolation and migration: insights from multiple mangrove taxa. Natl. Sci. Rev., 6, 275–288.

He, Z., Xu, S., Zhang, Z., Guo, W., Lyu, H., Zhong, C., et al. (2020) Convergent adaptation of the genomes of woody plants at the land–sea interface. Natl. Sci. Rev., 7, 978–993.

Hellsten, U., Wright, K.M., Jenkins, J., Shu, S., Yuan, Y., Wessler, S.R., et al. (2013) Fine-scale variation in meiotic recombination in Mimulus inferred from population shotgun sequencing. Proc. Natl. Acad. Sci., 110, 19478–19482.

Huang, S., Ding, J., Deng, D., Tang, W., Sun, Honghe, Liu, D., et al. (2013) Draft genome of the kiwifruit Actinidia chinensis. Nat. Commun., 4, 2640.

Jaillon, O., Aury, J.M., Noel, B., Policriti, A., Clepet, C., Casagrande, A., et al. (2007) The grapevine genome sequence suggests ancestral hexaploidization in major angiosperm phyla. Nature, 449, 463–467.

Ji, H., Pardo, J.M., Batelli, G., Van Oosten, M.J., Bressan, R.A., and Li, X. (2013) The Salt Overly Sensitive (SOS) Pathway: Established and Emerging Roles. Mol. Plant, 6, 275–286.

Jiao, Y., Leebens-Mack, J., Ayyampalayam, S., Bowers, J.E., McKain, M.R., McNeal, J., et al. (2012) A genome triplication associated with early diversification of the core eudicots. Genome Biol., 13, R3.

Katoh, K. (2002) MAFFT: a novel method for rapid multiple sequence alignment based on fast Fourier transform. Nucleic Acids Res., 30, 3059–3066.

Katoh, K. and Standley, D.M. (2013) MAFFT Multiple Sequence Alignment Software Version 7: Improvements in Performance and Usability. Mol. Biol. Evol., 30, 772–780.

Kim, D., Pertea, G., Trapnell, C., Pimentel, H., Kelley, R., and Salzberg, S.L. (2013) TopHat2: accurate alignment of transcriptomes in the presence of insertions, deletions and gene fusions. Genome Biol., 14, R36.

Kimura, M. (1980) A simple method for estimating evolutionary rates of base substitutions through comparative studies of nucleotide sequences. J. Mol. Evol., 16, 111–120.

Koren, S., Walenz, B.P., Berlin, K., Miller, J.R., Bergman, N.H., and Phillippy, A.M. (2017) Canu: scalable and accurate long-read assembly via adaptive k -mer weighting and repeat separation. Genome Res., 27, 722–736.

Kozlov, A.M., Darriba, D., Flouri, T., Morel, B., and Stamatakis, A. (2019) RAxML-NG: a fast, scalable and user-friendly tool for maximum likelihood phylogenetic inference. Bioinformatics, 35, 4453–4455.

Krzywinski, M., Schein, J., Birol, I., Connors, J., Gascoyne, R., Horsman, D., et al. (2009) Circos: An information aesthetic for comparative genomics. Genome Res., 19, 1639–1645.

Lamesch, P., Berardini, T.Z., Li, D., Swarbreck, D., Wilks, C., Sasidharan, R., et al. (2012) The Arabidopsis Information Resource (TAIR): improved gene annotation and new tools. Nucleic Acids Res., 40, D1202–D1210.

Langmead, B. and Salzberg, S.L. (2012) Fast gapped-read alignment with Bowtie 2. Nat. Methods, 9, 357–359.

Larsson, A. (2014) AliView: a fast and lightweight alignment viewer and editor for large datasets. Bioinformatics, 30, 3276–3278.

Li, H. (2018) Minimap2: pairwise alignment for nucleotide sequences. Bioinformatics, 34, 3094–3100.

Li, H. and Durbin, R. (2009) Fast and accurate short read alignment with Burrows-Wheeler transform. Bioinformatics, 25, 1754–1760.

Liang, S., Zhou, R., Dong, S., and Shi, S. (2008) Adaptation to salinity in mangroves: Implication on the evolution of salt-tolerance. Sci. Bull., 53, 1708–1715.

Liu, B., Shi, Y., Yuan, J., Hu, X., Zhang, H., Li, N., et al. (2013) Estimation of genomic characteristics by analyzing k-mer frequency in de novo genome projects. arXiv:1308.2012v2.

Lyu, H., He, Z., Wu, C., and Shi, S. (2018) Convergent adaptive evolution in marginal environments: unloading transposable elements as a common strategy among mangrove genomes. New Phytol., 217, 428–438.

Ma, T., Wang, Junyi, Zhou, G., Yue, Z., Hu, Q., Chen, Y., et al. (2013) Genomic insights into salt adaptation in a desert poplar. Nat. Commun., 4, 2797.

Majoros, W.H., Pertea, M., and Salzberg, S.L. (2004) TigrScan and GlimmerHMM: two open source ab initio eukaryotic gene-finders. Bioinformatics, 20, 2878–2879.

Mendes, F.K., Vanderpool, D., Fulton, B., and Hahn, M.W. (2020) CAFE 5 models variation in evolutionary rates among gene families. Bioinformatics, 10.1093/bioinformatics/btaa1022.

Morris, J.L., Puttick, M.N., Clark, J.W., Edwards, D., Kenrick, P., Pressel, S., et al. (2018) The timescale of early land plant evolution. Proc. Natl. Acad. Sci., 115, E2274–E2283.

Munns, R. and Tester, M. (2008) Mechanisms of Salinity Tolerance. Annu. Rev. Plant Biol., 59, 651–681.

Nakabayashi, K., Bartsch, M., Xiang, Y., Miatton, E., Pellengahr, S., Yano, R., et al. (2012) The Time Required for Dormancy Release in Arabidopsis Is Determined by DELAY OF GERMINATION1 Protein Levels in Freshly Harvested Seeds. Plant Cell, 24, 2826–2838.

Nishimura, N., Tsuchiya, W., Moresco, J.J., Hayashi, Y., Satoh, K., Kaiwa, N., et al. (2018) Control of seed dormancy and germination by DOG1-AHG1 PP2C phosphatase complex via binding to heme. Nat. Commun., 9, 2132.

Nonogaki, H. (2019) Seed germination and dormancy: The classic story, new puzzles, and evolution. J. Integr. Plant Biol., 61, 541–563.

Nowak, M.D., Russo, G., Schlapbach, R., Huu, C.N., Lenhard, M., and Conti, E. (2015) The draft genome of Primula veris yields insights into the molecular basis of heterostyly. Genome Biol., 16, 12.

Ouyang, S., Zhu, W., Hamilton, J., Lin, H., Campbell, M., Childs, K., et al. (2007) The TIGR Rice Genome Annotation Resource: improvements and new features. Nucleic Acids Res., 35, D883–D887.

Parida, A.K., Das, A.B., Sanada, Y., and Mohanty, P. (2004) Effects of salinity on biochemical components of the mangrove, Aegiceras corniculatum. Aquat. Bot., 80, 77–87.

Parida, A.K. and Jha, B. (2010) Salt tolerance mechanisms in mangroves: a review. Trees, 24, 199–217.

Van De Peer, Y., Ashman, T.-L., Soltis, P.S., and Soltis, D.E. (2020) Polyploidy: an Evolutionary and Ecological Force in Stressful Times. Plant Cell, 10.1093/plcell/koaa015.

Van de Peer, Y., Mizrachi, E., and Marchal, K. (2017) The evolutionary significance of polyploidy. Nat. Rev. Genet., 18, 411–424.

Qiao, H., Zhou, X., Su, W., Zhao, X., Jin, P., He, S., et al. (2020) The genomic and transcriptomic foundations of viviparous seed development in mangroves. bioRxiv, 10.1101/2020.10.19.346163.

Reis, M. Dos and Yang, Z. (2011) Approximate Likelihood Calculation on a Phylogeny for Bayesian Estimation of Divergence Times. Mol. Biol. Evol., 28, 2161–2172.

Roach, M.J., Schmidt, S.A., and Borneman, A.R. (2018) Purge Haplotigs: allelic contig reassignment for third-gen diploid genome assemblies. BMC Bioinformatics, 19, 460.

Seki, M., Narusaka, M., Ishida, J., Nanjo, T., Fujita, M., Oono, Y., et al. (2002) Monitoring the expression profiles of 7000 Arabidopsis genes under drought, cold and high-salinity stresses using a full-length cDNA microarray. Plant J., 31, 279–292.

Seppey, M., Manni, M., and Zdobnov, E.M. (2019) BUSCO: Assessing Genome Assembly and Annotation Completeness. Methods in Molecular Biology, 227–245.

Shannon, P., Markiel, A., Ozier, O., Baliga, N.S., Wang, J.T., Ramage, D., et al. (2003) Cytoscape: A Software Environment for Integrated Models of Biomolecular Interaction Networks. Genome Res., 13, 2498–2504.

Shi, S., Huang, Y., Zeng, K., Tan, F., He, H., Huang, J., and Fu, Y. (2005) Molecular phylogenetic analysis of mangroves: independent evolutionary origins of vivipary and salt secretion. Mol. Phylogenet. Evol., 34, 159–166.

Shi, T., Rahmani, R.S., Gugger, P.F., Wang, M., Li, H., Zhang, Y., et al. (2020) Distinct Expression and Methylation Patterns for Genes with Different Fates following a Single Whole-Genome Duplication in Flowering Plants. Mol. Biol. Evol., 37, 2394–2413.

Stanke, M., Keller, O., Gunduz, I., Hayes, A., Waack, S., and Morgenstern, B. (2006) AUGUSTUS: ab initio prediction of alternative transcripts. Nucleic Acids Res., 34, W435–W439.

Sugimoto, K., Takeuchi, Y., Ebana, K., Miyao, A., Hirochika, H., Hara, N., et al. (2010) Molecular cloning of Sdr4, a regulator involved in seed dormancy and domestication of rice. Proc. Natl. Acad. Sci., 107, 5792–5797.

Suo, Y., Sun, P., Cheng, H., Han, W., Diao, S., Li, H., et al. (2020) A high-quality chromosomal genome assembly of Diospyros oleifera Cheng. Gigascience, 9, 1–10.

Suyama, M., Torrents, D., and Bork, P. (2006) PAL2NAL: robust conversion of protein sequence alignments into the corresponding codon alignments. Nucleic Acids Res., 34, W609–W612.

Tarailo-Graovac, M. and Chen, N. (2009) Using RepeatMasker to Identify Repetitive Elements in Genomic Sequences. Curr. Protoc. Bioinforma., 25, 1–14.

Tomlinson, P.B. (2016) The botany of mangroves. 2nd ed. Cambridge University Press.

Tomlinson, P.B. and Cox, P.A. (2000) Systematic and functional anatomy of seedlings in mangrove Rhizophoraceae: vivipary explained? Bot. J. Linn. Soc., 134, 215–231.

Trapnell, C., Roberts, A., Goff, L., Pertea, G., Kim, D., Kelley, D.R., et al. (2012) Differential gene and transcript expression analysis of RNA-seq experiments with TopHat and Cufflinks. Nat. Protoc., 7, 562–578.

Vaser, R., Sović, I., Nagarajan, N., and Šikić, M. (2017) Fast and accurate de novo genome assembly from long uncorrected reads. Genome Res., 27, 737–746.

Walker, B.J., Abeel, T., Shea, T., Priest, M., Abouelliel, A., Sakthikumar, S., et al. (2014) Pilon: An Integrated Tool for Comprehensive Microbial Variant Detection and Genome Assembly Improvement. PLoS One, 9, e112963.

Wang, D., Zhang, Y., Zhang, Z., Zhu, J., and Yu, J. (2010) KaKs_Calculator 2.0: A Toolkit Incorporating Gamma-Series Methods and Sliding Window Strategies. Genomics. Proteomics Bioinformatics, 8, 77–80.

Wang, Y., Tang, H., DeBarry, J.D., Tan, X., Li, J., Wang, X., et al. (2012) MCScanX: a toolkit for detection and evolutionary analysis of gene synteny and collinearity. Nucleic Acids Res., 40, e49.

Wu, S., Han, B., and Jiao, Y. (2020) Genetic Contribution of Paleopolyploidy to Adaptive Evolution in Angiosperms. Mol. Plant, 13, 59–71.

Xia, E., Li, F., Tong, W., Li, P., Wu, Q., Zhao, H., et al. (2019) Tea Plant Information Archive: a comprehensive genomics and bioinformatics platform for tea plant. Plant Biotechnol. J., 17, 1938–1953.

Xiao, C.-L., Chen, Y., Xie, S.-Q., Chen, K.-N., Wang, Y., Han, Y., et al. (2017) MECAT: fast mapping, error correction, and de novo assembly for single-molecule sequencing reads. Nat. Methods, 14, 1072–1074.

Xu, S., He, Z., Zhang, Z., Guo, Z., Guo, W., Lyu, H., et al. (2017) The origin, diversification and adaptation of a major mangrove clade (Rhizophoreae) revealed by whole-genome sequencing. Natl. Sci. Rev., 4, 721–734.

Xu, Z. and Wang, H. (2007) LTR_FINDER: an efficient tool for the prediction of full-length LTR retrotransposons. Nucleic Acids Res., 35, W265–W268.

Yang, F.-S., Nie, S., Liu, H., Shi, T.-L., Tian, X.-C., Zhou, S.-S., et al. (2020) Chromosome-level genome assembly of a parent species of widely cultivated azaleas. Nat. Commun., 11, 5269.

Yang, Z. (2007) PAML 4: Phylogenetic Analysis by Maximum Likelihood. Mol. Biol. Evol., 24, 1586–1591.

Yang, Z., Wang, C., Xue, Y., Liu, X., Chen, S., Song, C., et al. (2019) Calcium-activated 14-3-3 proteins as a molecular switch in salt stress tolerance. Nat. Commun., 10, 1199.

Yu, G., Smith, D.K., Zhu, H., Guan, Y., and Lam, T.T. (2017) ggtree: an package for visualization and annotation of phylogenetic trees with their covariates and other associated data. Methods Ecol. Evol., 8, 28–36.

Yuan, F., Leng, B., and Wang, B. (2016) Progress in Studying Salt Secretion from the Salt Glands in Recretohalophytes: How Do Plants Secrete Salt? Front. Plant Sci., 7, 977.

Zhang, L., Xu, P., Cai, Y., Ma, L., Li, Shifeng, Li, Shufa, et al. (2017) The draft genome assembly of Rhododendron delavayi Franch. var. delavayi. Gigascience, 6, 1–11.

Zheng, W., Wang, W., and Lin, P. (1999) Dynamics of element contents during the development of hypocotyles and leaves of certain mangrove species. J. Exp. Mar. Bio. Ecol., 233, 247–257.

